# A proteolytic complex targets multiple cell wall hydrolases in *Pseudomonas aeruginosa*

**DOI:** 10.1101/260620

**Authors:** Disha Srivastava, Jin Seo, Binayak Rimal, Sung Joon Kim, Andrew J. Darwin

**Affiliations:** Department of Microbiology, New York University School of Medicine, New York, New York, United States; Institute of Biomedical Studies, Baylor University, Waco, Texas, United States; Department of Chemistry and Biochemistry, Baylor University, Waco, Texas, United States

## Abstract

Carboxy-terminal processing proteases (CTPs) occur in all three domains of life. In bacteria some of them have been associated with virulence. However, the precise roles of bacterial CTPs are poorly understood and few direct proteolytic substrates have been identified. One bacterial CTP is the CtpA protease of *Pseudomonas aeruginosa*, which is required for type III secretion system function, and for virulence in a mouse model of acute pneumonia. Here, we have investigated the function of CtpA in *P. aeruginosa* and identified some of the proteins it cleaves. We discovered that CtpA forms a complex with a previously uncharacterized protein, which we have named LbcA (<u>l</u>ipoprotein <u>b</u>inding partner of <u>C</u>tpA). LbcA is required for CtpA activity *in vivo* and promotes its activity *in vitro*. We have also identified four proteolytic substrates of CtpA, all of which are uncharacterized proteins predicted to cleave the peptide cross-links within peptidoglycan. Consistent with this, a *ctpA* null mutant was found to have fewer peptidoglycan cross-links than the wild type and grew slowly in salt-free medium. Intriguingly, the accumulation of just one of the CtpA substrates was required for some Δ*ctpA* mutant phenotypes, including the defective T3SS. We propose that LbcA•CtpA is a proteolytic complex in the *P. aeruginosa* cell envelope, which controls the activity of several peptidoglycan cross-link hydrolases by degrading them. Furthermore, based on these and other findings we suggest that many bacterial CTPs might be similarly controlled by partner proteins as part of a widespread mechanism to control peptidoglycan hydrolase activity.

**IMPORTANCE:** Bacterial carboxy-terminal processing proteases (CTPs) are widely conserved and have been associated with the virulence of several species. However, their roles are poorly understood and few direct substrates have been identified in any species. *Pseudomonas aeruginosa* is an important human pathogen in which one CTP, known as CtpA, is required for type III secretion system function, and for virulence. This work provides an important advance by showing that CtpA works with a previously uncharacterized binding partner to degrade four substrates. These substrates are all predicted to hydrolyze peptidoglycan cross-links, suggesting that the CtpA complex is an important control mechanism for peptidoglycan hydrolysis. This is likely to emerge as a widespread mechanism used by diverse bacteria to control some of their peptidoglycan hydrolases. This is significant, given the links between CTPs and virulence in several pathogens, and the importance of peptidoglycan remodeling to almost all bacterial cells.

## INTRODUCTION

*Pseudomonas aeruginosa* is a Gram-negative bacterium that accounts for over 10% of healthcare associated infections with an identifiable cause (1). In acute infections, *P. aeruginosa* utilizes a type III secretion system (T3SS) to export proteins into host cells, where they interfere with signaling, immune function, and cause cytotoxicity (2). In chronic infections, *P. aeruginosa* forms biofilms resilient to clearance by antibiotics and the immune response (3, 4). The emergence of antibiotic resistant strains is increasing, making this pathogen a top priority for the discovery of new therapeutic targets (5).

Proteases are important virulence factors for many pathogens, including *P. aeruginosa* (6). One intriguing family is the <u>c</u>arboxy-<u>t</u>erminal <u>p</u>rocessing proteases (CTPs), found in all three domains of life. CTPs belong to the S41 family of serine proteases with a serine/lysine catalytic dyad (7). They are believed to target substrates close to their C-terminus (8, 9). A well described CTP occurs in plant chloroplasts and cyanobacteria, where it cleaves a component of the photosystem II reaction center to activate it (10, 11). Gram-negative bacterial CTPs are found exclusively in the periplasmic compartment and have been implicated in cleaving proteins associated with cell envelope function (8, 12). Some CTPs have been associated with virulence, but mostly by poorly defined mechanisms (13–19). Indeed, despite their widespread conservation, our knowledge of how CTPs function, and what they cleave, is limited.

*Escherichia coli* has one CTP, known as Prc or Tsp (8, 9). *prc* null mutants have altered cell morphology, increased drug and low osmotic sensitivity, and altered virulence (20, 21). Prc was proposed to process penicillin binding protein 3 (FtsI), an essential peptidoglycan transpeptidase required for cell division (8). More recently another Prc substrate was discovered, the peptidoglycan hydrolase MepS, which cleaves peptide cross-links between the glycan chains of peptidoglycan (22, 23). These substrates suggest that Prc plays a role in controlling different aspects of peptidoglycan metabolism. However, Prc has also been shown to cleave incorrectly folded polypeptides in the periplasm, suggesting that it has a broader role in protein quality control (9, 24, 25). *P. aeruginosa* has a homolog of *E. coli* Prc, with the two proteins sharing over 40% amino acid identity and belonging to the CTP-1 subfamily (7). *P. aeruginosa* Prc has been proposed to contribute to degradation of the antisigma factor MucA, which induces the AlgT/U regulon and subsequent production of the exopolysaccharide, alginate (26–28).

Unlike *E. coli, P. aeruginosa* has a second CTP, CtpA, which is smaller than Prc and in a different subfamily (the CTP-3 subfamily; 18, 29). CtpA is required for T3SS function, and a *ctpA* null mutant is less cytotoxic to cultured cells and attenuated in a mouse model of acute pneumonia (18). We also reported that a Δ*ctpA* mutant is sensitive to mislocalized pore forming secretin proteins, has increased resistance to two cationic surfactants, has an altered cell envelope when viewed by electron microscopy, and that CtpA overproduction induces an extracytoplasmic function sigma factor regulon (18). Furthermore, others found that a *ctpA* transposon insertion mutant has reduced swarming motility (30). All of this suggests that CtpA has a significant impact on the cell envelope. Here, we report that CtpA forms a complex with a protein we have named LbcA (<u>l</u>ipoprotein <u>b</u>inding partner of <u>C</u>tpA). LbcA is required for CtpA activity *in vivo* and promotes its activity *in vitro*. We have also identified four substrates of CtpA, all of which are uncharacterized proteins predicted to cleave the peptide cross-links within peptidoglycan. The accumulation of one of these substrates appears to be needed for some of the most significant phenotypes of a Δ*ctpA* mutant, including the defective T3SS. We argue that many bacterial CTPs might be controlled by partner proteins as part of a widespread mechanism to control peptidoglycan hydrolase activity.

## RESULTS

### Identification of a CtpA binding partner

We modified the chromosomal *ctpA* gene to encode CtpA-FLAG-His6, in strains encoding wild type CtpA or CtpA-S302A (catalytic serine mutated to alanine). The rationale was that CtpA-S302A might trap substrates after tandem affinity purification. These experiments were done with or without *in vivo* formaldehyde cross-linking prior to protein purification, but the results were similar and only an experiment without cross-linking is shown (Fig. 1A).

**FIG 1.**
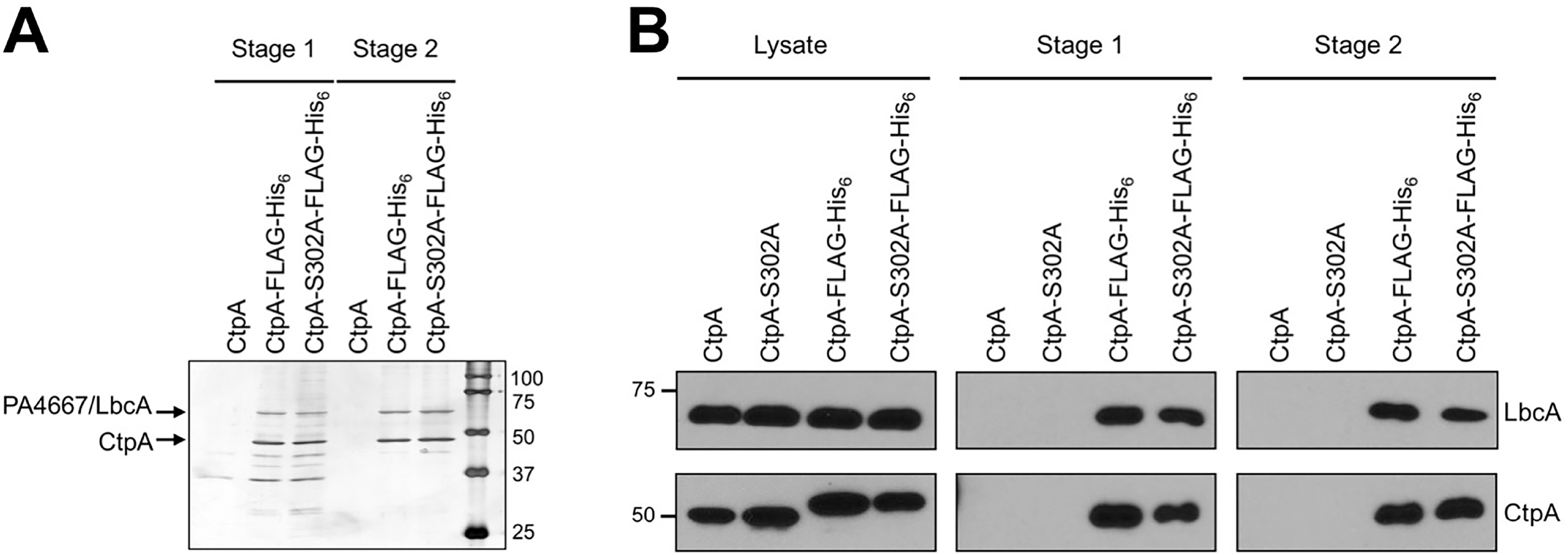
Identification of the CtpA interaction partner PA4667/LbcA. (A) Tandem affinity tag purification. Proteins were purified from detergent solubilized lysates of the wild type strain (CtpA), or derivatives encoding CtpA-FLAG-His_6_ or the proteolytically inactive CtpA-S302A-FLAG-His_6_. Purification was done with nickel agarose (Stage 1) followed by anti-FLAG M2 agarose resin (Stage 2). Samples were separated on a 12.5% SDS polyacrylamide gel, which was stained with silver. Molecular-mass-marker proteins (kDa) are labeled on the right-hand side. (B) Immunoblot analysis of detergent solubilized lysate, and Stage 1 and 2 purified samples, prepared as in panel A. LbcA and CtpA were detected with polyclonal antisera. Strains in which CtpA or CtpA-S302A were not tagged with FLAG-His_6_ served as negative controls. Approximate positions of molecular-mass-marker proteins (kDa) are indicated on the left-hand side.

Both versions of CtpA co-purified with abundant amounts of a protein running between 50 and 75 kDa markers in SDS-PAGE (Fig. 1A). Mass spectrometry identified it as PA4667 (PAO1 strain designation). A polyclonal antiserum raised against PA4667 confirmed its co-purification with epitope tagged CtpA or CtpA-S302A (Fig 1B). PA4667 is a predicted 63 kDa outer membrane lipoprotein with eleven tetratricopeptide (TPR) repeats. The TPR motif is a degenerate 34 amino acid sequence that mediates protein-protein interactions (31). PA4667 does not appear to be a CtpA substrate because its level is similar in *ctpA*^+^ and Δ*ctpA* strains (see below). Therefore, we will refer to PA4667 as LbcA, (<u>l</u>ipoprotein <u>b</u>inding partner of <u>C</u>tpA).

### Δ*lbcA* and Δ*ctpA* mutants have common phenotypes

We hypothesized that LbcA might be required for CtpA function, but we could not construct Δ*lbcA* in frame deletion mutants. We could make mutants where *lbcA* was replaced by *aacC1*, encoding gentamycin resistance, but only if *aacC1* was in the same orientation as *lbcA. lbcA* is in an operon with the downstream essential genes, *lolB* and *ipk* (32). We speculate that the *aacC1* promoter helps to express *lolB-ipk*, and the inability to make a Δ*lbcA* in frame deletion might be due to reduced *lolB-ipk* expression.

A Δ*ctpA* mutant has a defective T3SS (18). Therefore, we compared the ability of *ctpA* and *lbcA* null mutants to export T3SS effector proteins ExoS and ExoT and found that both were defective (Fig. 2A). The Δ*lbcA::aacC1* mutant was complemented by an *lbcA*^+^ plasmid, and the Δ*ctpA* mutant was complemented by wild type CtpA but not by CtpA-S302A (Fig. 2A).

**FIG 2.**
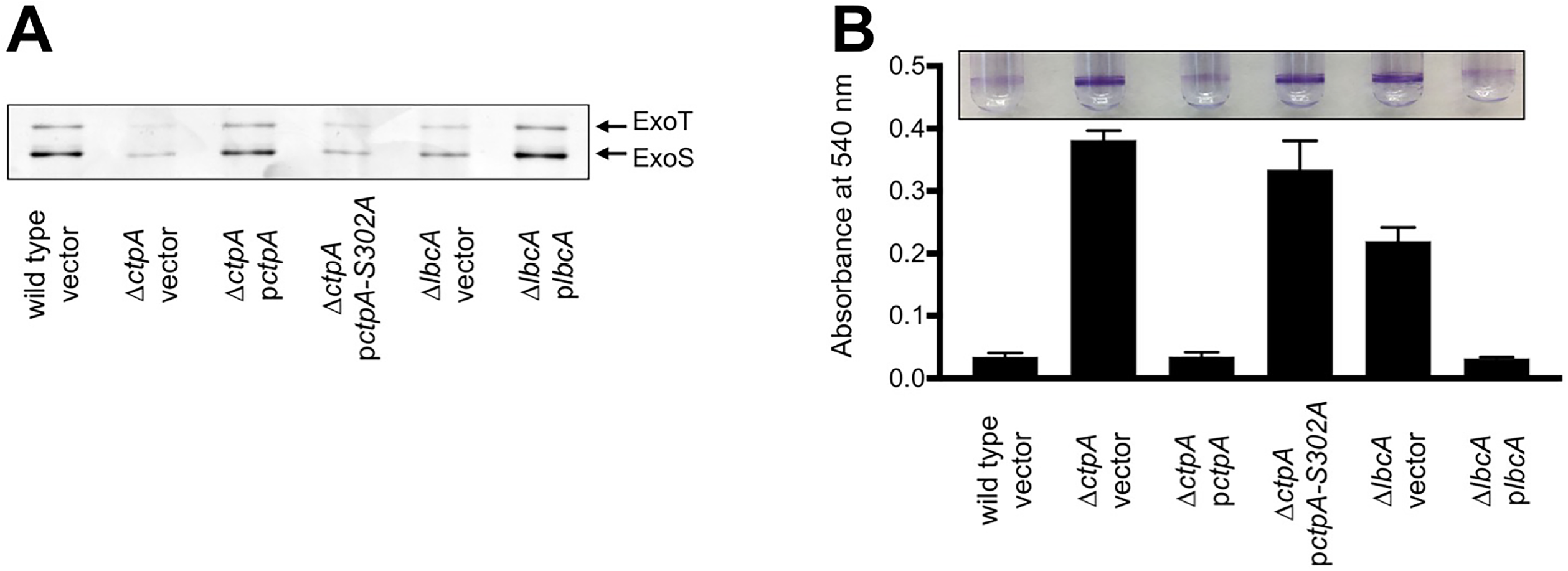
Δ*ctpA* and Δ*lbcA* mutants share common phenotypes. (A) T3SS function. Wild type, Δ*ctpA* or Δ*lbcA* strains contained the *tac* promoter expression plasmid pVLT35 (vector), or derivatives encoding CtpA, CtpA-S302A or LbcA as indicated. TSB growth medium contained 1mM EGTA to induce T3SS activity, and 10 μM IPTG to induce the *tac* promoter of the complementing plasmids. Cell free supernatants derived from equivalent amounts of cells were separated on a SDS-PAGE gel, which was stained with silver. The region with the abundant T3SS effectors ExoT and ExoS is shown. (B) Surface attachment. The same strains as in panel A were incubated in glass test tubes at 30°C without agitation for 8 hours. The growth medium contained 10μM IPTG to induce the *tac* promoter of the complementing plasmids. Bacterial cells attached to the glass at the air/liquid interface were stained with crystal violet (purple rings). The stain was dissolved in ethanol and quantified by measuring absorbance at 595 nm. All strains were tested in triplicate and error bars represent the standard deviation from the mean.

Recently, we discovered that a Δ*ctpA* mutant has a phenotype in a commonly used surface attachment assay (33). After an 8 hour incubation, the Δ*ctpA* mutant attached robustly whereas the wild type did not (Fig. 2B). We tested if the *lbcA* null mutant shares this phenotype, and it did (Fig. 2B). After longer incubation the wild type caught up so that attachment of wild type and mutant strains was indistinguishable (data not shown). This accelerated attachment phenotype was not caused by faster growth of the mutants in these assays (data not shown).

### Evidence that LbcA recruits CtpA to the outer membrane

LbcA is a predicted outer membrane lipoprotein, so it should tether CtpA to the membrane. However, we and others concluded that CtpA is a soluble periplasmic protein because it was released by osmotic shock (18, 29). Even so, some CtpA was not released in our experiment (18). Indeed, we have found that a significant amount of CtpA is always retained, and this is more pronounced when cultures are harvested at a lower optical density (OD) than before (OD 600nm of 1 instead of 1.5). Therefore, we osmotically shocked *lbcA*^+^ and Δ*lbcA* strains grown to OD 600 nm of 1. In the *lbcA*^+^ strain, most CtpA was retained, whereas a soluble periplasmic control protein was released (Fig. 3A). However, in an Δ*lbcA* mutant, CtpA was released. This supports the hypothesis that LbcA influences CtpA localization.

**FIG 3.**
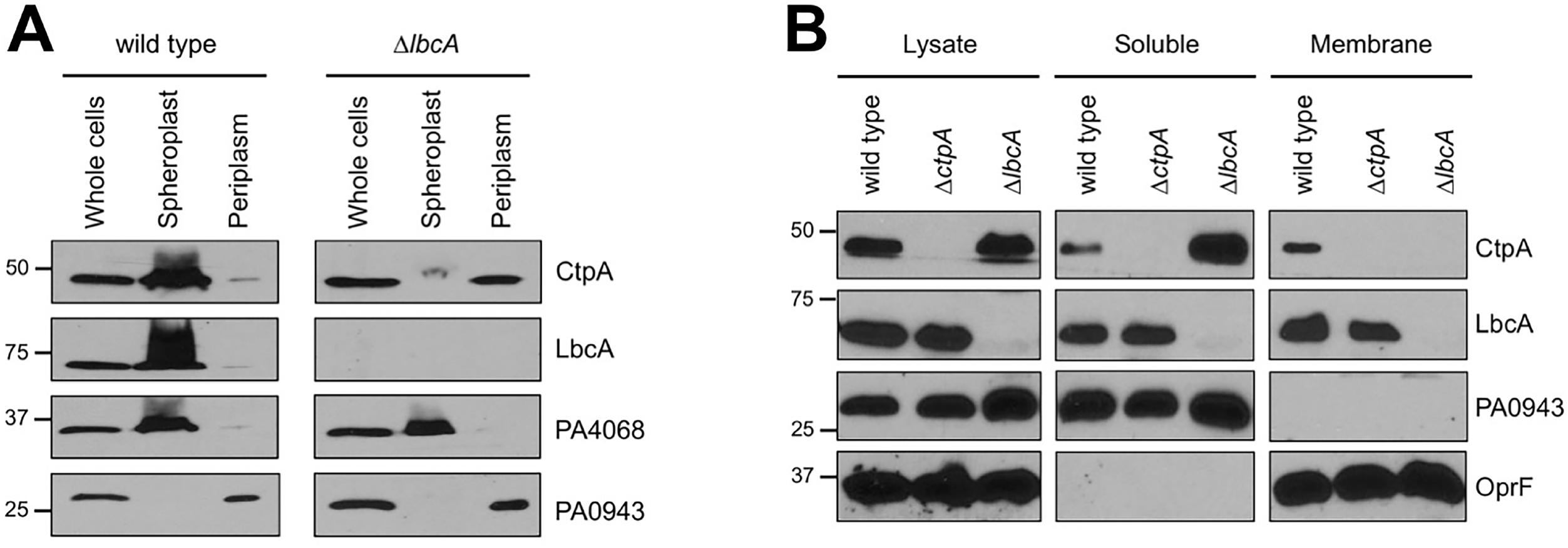
LbcA promotes retention of CtpA in spheroplast and membrane fractions. (A) Osmotic shock fractionation. Total cell lysate (whole cells), as well as spheroplast and periplasm fractions generated by osmotic shock, were separated by SDS-PAGE, transferred to nitrocellulose and proteins detected with polyclonal antisera. PA0943 is a soluble periplasmic protein control, and PA4068 is a cytoplasmic spheroplast-retention control (18). (B) Fractionation of lysed cells. Cells were lysed by two passages through a French pressure cell and separated into soluble and insoluble (Membrane) fractions by ultracentrifugation. Samples derived from equivalent amounts of cells were separated by SDS PAGE, transferred to nitrocellulose, and proteins detected with polyclonal antisera. PA0943 is a soluble protein control and OprF is an insoluble outer membrane control.

We also separated cells into soluble and insoluble fractions by lysis and ultracentrifugation. Interestingly, LbcA was found in the soluble and insoluble fractions (Fig. 3B). We do not know if this is physiologically significant, or an artifact resulting from LbcA being solubilized during sample processing. Regardless, in an *lbcA*^+^ strain, CtpA fractionated identically to LbcA, whereas in an Δ*lbcA* mutant CtpA was only in the soluble fraction (Fig. 3B). This is also consistent with an LbcA•CtpA interaction tethering some CtpA to the outer membrane.

### Isolation of a putative CtpA substrate

To find substrates of the LbcA•CtpA complex, we modified our tandem affinity tag approach by putting the His_6_ tag on CtpA or CtpA-S302A, and the FLAG tag on LbcA. Strains were grown in T3SS inducing conditions and treated with formaldehyde to cross-link associated proteins. Proteins were purified using nickel agarose followed by anti-FLAG affinity gel and analyzed by SDS-PAGE (Fig. 4A). We also analyzed two independent purifications from each strain by mass spectrometry. 101 proteins were present in both LbcA•CtpA-S302A samples, but neither LbcA•CtpA sample (Fig. 4B and Supplemental Table 1). We focused on the most abundant according to peptide spectrum matches (PSMs), the uncharacterized PA0667 (Fig. 4B). PA0667 has a Sec-dependent signal sequence, a LysM peptidoglycan-binding domain, and a LytM/M23 peptidase domain. It is 35% identical to *E. coli* MepM, a DD-endopeptidase that cleaves d-Ala-meso-DAP peptide peptidoglycan cross-links (23). Therefore, we will refer to PA0667 as MepM.

**FIG 4.**
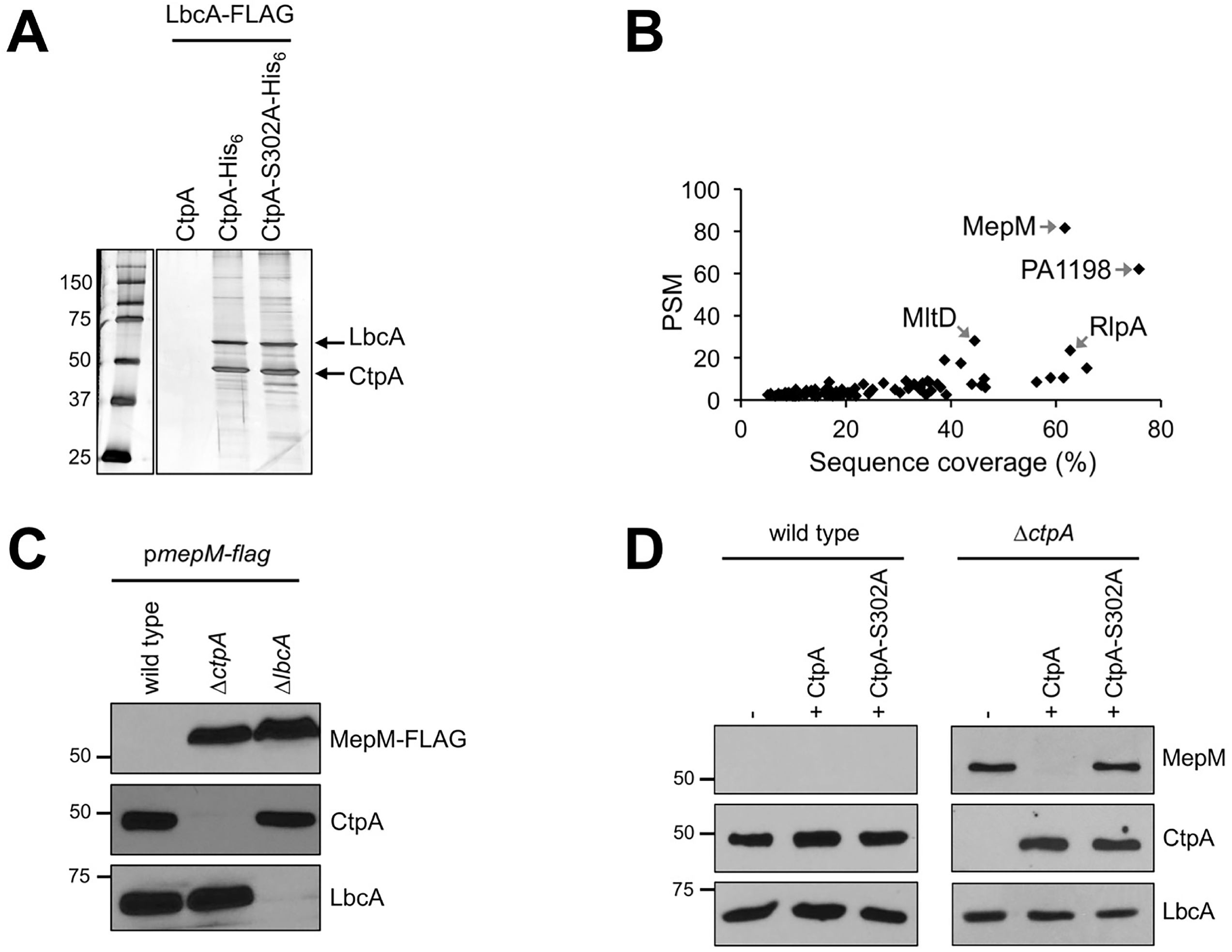
Identification of MepM as a putative substrate of the LbcA•CtpA proteolytic complex. (A) Tandem affinity tag purification. Proteins were purified from detergent solubilized lysates of Δ*ctpA* strains. All strains had a plasmid encoding LbcA-FLAG as well as a second plasmid encoding untagged CtpA (negative control), CtpA-His_6_ or the proteolytically inactive CtpA-S302A-His_6_. Purification was done with nickel agarose followed by anti-FLAG M2 agarose resin. Samples were separated on a 12.5% SDS polyacrylamide gel, which was stained with silver. Molecular-mass-marker proteins (kDa) are labeled on the left-hand side. An irrelevant region of the gel between the marker and sample lanes has been removed. (B) Scatter plot of proteins identified by mass spectrometry after purification from strains encoding CtpA-S302A-His_6_. Data are the average peptide spectrum matches (PSM) and sequence coverage from duplicate purifications from strains with CtpA-S302A-His6. Each point represents a different protein. All proteins plotted were detected in both of the LbcA•CtpA-S302A-His_6_ purifications but neither of duplicate LbcA•CtpA-His_6_ purifications. (C) Immunoblot analysis of equivalent amounts of whole cell lysates of wild type, Δ*ctpA* and Δ*lbcA* strains. All strains contained an arabinose-inducible expression plasmid encoding MepM-FLAG and were grown in media containing 0.2% (w/v) arabinose. Approximate positions of molecular-mass-marker proteins (kDa) are indicated on the left-hand side. MepM-FLAG was detected with anti-FLAG monoclonal antibodies and CtpA and LbcA were detected with polyclonal antisera. (D) Detection of endogenous MepM. Immunoblot analysis of equivalent amounts of whole cell lysates of wild type and Δ*ctpA* strains. Strains contained the *tac* promoter expression plasmid pVLT35 (-), or derivatives encoding CtpA or CtpA-S302A as indicated. Approximate positions of molecular-mass-marker proteins (kDa) are indicated on the left-hand side. MepM, CtpA and LbcA were detected with polyclonal antisera.

### Evidence that MepM is a substrate of the LbcA•CtpA complex *in vivo*

To test if MepM might be an LbcA•CtpA substrate, we constructed a plasmid encoding arabinose inducible MepM-FLAG. After growth in the presence of arabinose the MepM-FLAG protein was undetectable in the wild type strain, whereas it was abundant in Δ*ctpA* and Δ*lbcA* mutants (Fig. 4C). To analyze endogenous MepM, we raised a polyclonal antiserum against it. This antiserum did not detect MepM in the wild type strain, but once again it revealed MepM accumulated in a Δ*ctpA* mutant (Fig. 4D). This accumulation was reversed by a plasmid encoding wild type CtpA, but not by CtpA-S302A (Fig 4D). These data suggest that MepM is cleaved by CtpA *in vivo*.

### Deletion of *mepM* suppresses Δ*ctpA* mutant phenotypes

Δ*ctpA* phenotypes might be caused by MepM accumulation. If so, a Δ*mepM* mutation should suppress them. To test this, we introduced a *mepM* in frame deletion mutation into wild type and Δ*ctpA* strains and surveyed their phenotypes (Fig. 5).

**FIG 5.**
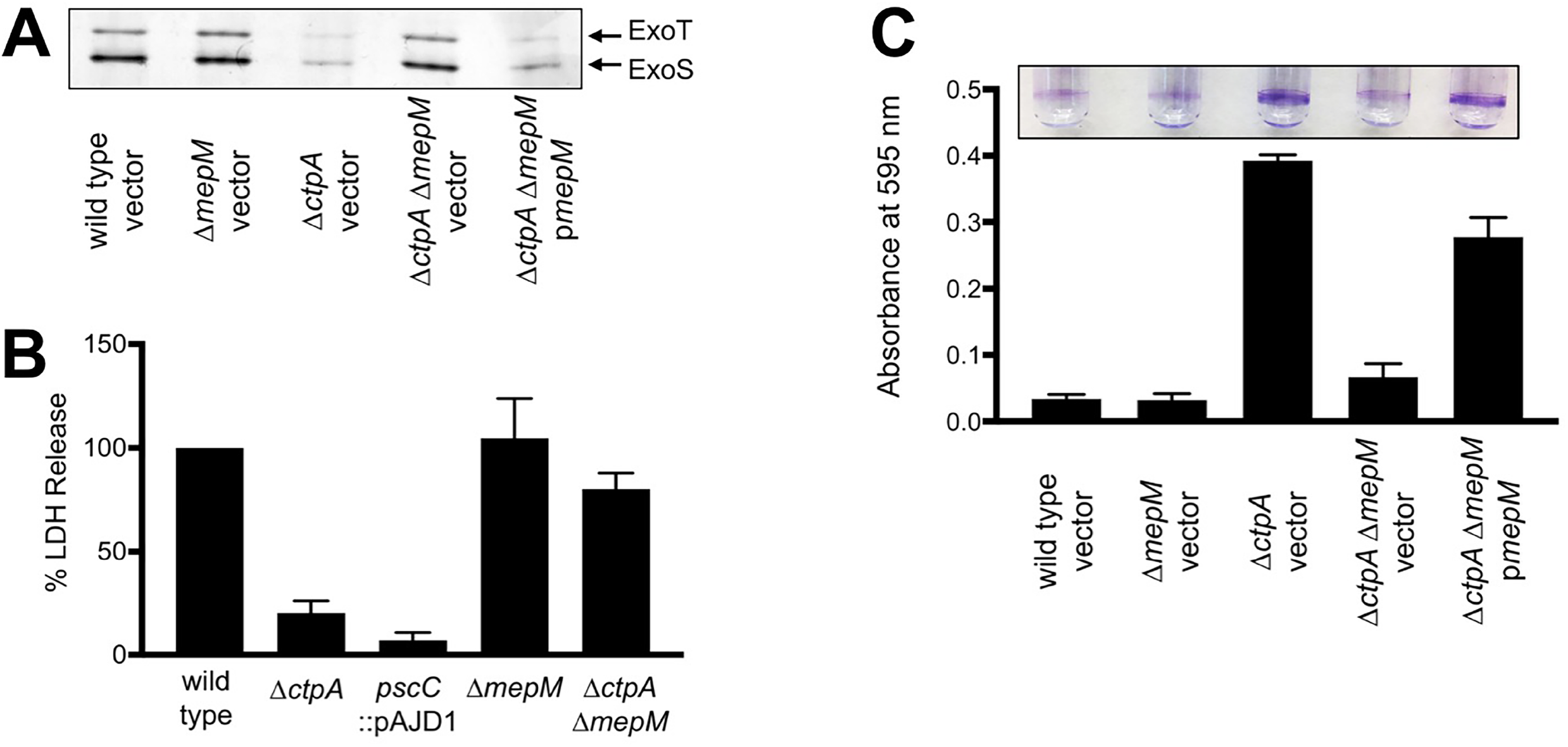
*mepM* null mutation suppresses Δ*ctpA* phenotypes. (A) T3SS function. Wild type, Δ*mepM*, Δ*ctpA* or Δ*ctpA* Δ*mepM* strains contained the *araBp* promoter expression plasmid pHERD20T (vector), or a derivative encoding MepM as indicated. TSB growth medium contained 1 mM EGTA to induce T3SS activity, and 0.02% arabinose to induce the *araB* promoter of pHERD20T. Cell free supernatants derived from equivalent amounts of cells were separated by SDS-PAGE gel, which was stained with silver. The region with the abundant T3SS effectors ExoT and ExoS is shown. (B) Cytotoxicity to CHO-K1 cells. Strains with the indicated genotypes were added to CHO-K1 cells at a multiplicity of infection of ~10 and incubated for 4 h. Cell-free supernatants were then analyzed for lactate dehydrogenase (LDH) content. The amount of LDH in the supernatant following incubation with the wild type strain was set to 100% and the values for the mutants are shown as the relative percentage. Data were averaged from three independent experiments with the error bars showing the standard deviation from the mean. (C) Surface attachment. The same strains from panel A were incubated in glass test tubes at 30°C without agitation for 8 hours. The growth medium contained 0.02% arabinose to induce the *araB* promoter of the complementing plasmids. Bacterial cells attached to the glass at the air liquid interface were stained with crystal violet (purple rings). The stain was dissolved in ethanol and quantified by measuring absorbance at 595nm. All strains were tested in triplicate and error bars represent the standard deviation from the mean.

First, we monitored T3SS function. The Δ*mepM* mutation alone did not affect ExoS/T export (Fig. 5A), whereas a Δ*ctpA* mutant exported lower amounts, as reported before (18). However, the Δ*ctpA* phenotype was suppressed by a Δ*mepM* mutation, with the ExoS/T secretion profiles of wild type and Δ*ctpA* Δ*mepM* strains being indistinguishable (Fig. 5A). Furthermore, *mepM* expression from a plasmid reduced T3SS function in the Δ*ctpA* Δ*mepM* mutant (Fig 5A). These data suggest that MepM accumulation was responsible for the defective T3SS of the Δ*ctpA* mutant.

We also investigated cytotoxicity towards CHO-K1 cells by measuring lactate dehydrogenase (LDH) release. This is an established model for studying the toxic effect of the *P. aeruginosa* T3SS on eukaryotic cells (18, 34). The cytotoxicity phenotypes of the Δ*ctpA*, Δ*mepM* and Δ*ctpA* Δ*mepM* strains were consistent with their T3SS phenotypes (Figs. 5A and B). Δ*mepM* did not affect cytotoxicity, Δ*ctpA* reduced it, and this reduction was suppressed in the Δ*ctpA* Δ*mepM* double mutant (Fig. 5B).

Finally, the Δ*mepM* mutation also suppressed the accelerated surface attachment phenotype of a Δ*ctpA* mutant, and this suppression was reversed by a *mepM*^+^ plasmid (Fig. 5C). Together, all these findings suggest that many phenotypes of a Δ*ctpA* mutant require MepM. As expected, a Δ*mepM* mutation also suppressed these phenotypes of a Δ*lbcA* mutant (data not shown). Therefore, we hypothesized that the phenotypes of LbcA•CtpA-defective strains are the result of increased hydrolysis of peptidoglycan cross-links, which we tested next.

### Peptidoglycan cross-linking is reduced in a Δ*ctpA* mutant

The peptidoglycan compositions of wild type and Δ*ctpA* strains were characterized by mass spectrometry. The repeat unit consists of the disaccharide Glc*N*Ac-Mur*N*Ac with a pentapeptide stem l-Ala-d-iso-Gln-*m*-Dap-d-Ala-d-Ala attached to the lactic moiety of Mur*N*Ac (Fig. 6A). The disaccharides are joined by β_1-4_ glycosidic linkages to form the glycan chain. The glycan chains are cross-linked via a peptide bond between the sidechain of *m*-DAP in an acceptor stem of one repeat unit, to the carbonyl carbon of a penultimate d-Ala in a donor stem of a neighboring glycan chain. Cell walls were digested with mutanolysin, which cleaves the β_1-4_ glycosidic linkage between Mur*N*Ac and Glc*N*Ac, resulting in muropeptide fragments with intact cross-links. Muropeptide ions were identified by matching the observed m/z values to the corresponding entries in a muropeptide library generated *in silico* using an in-house program. Forty-seven unique muropeptide ion species from the wild type, and thirty-three from the Δ*ctpA* mutant were selected for the analysis. Muropeptide fragments were quantified by integrating the extracted ion chromatogram (XIC) of the selected muropeptide ion (35–37). The normalized summed integrals of muropeptides from the XIC ion current were categorized based on the cross-link number per muropeptide fragment (Fig. 6B). Muropeptide fragments larger than trimers were not observed, and the most abundant muropeptides were dimers. Representative spectra of monomer, dimer, and trimer from the wild type are shown in Figure 6, and the chemical structure and calculated masses are in Supplementary Figure S1.

**FIG 6.**
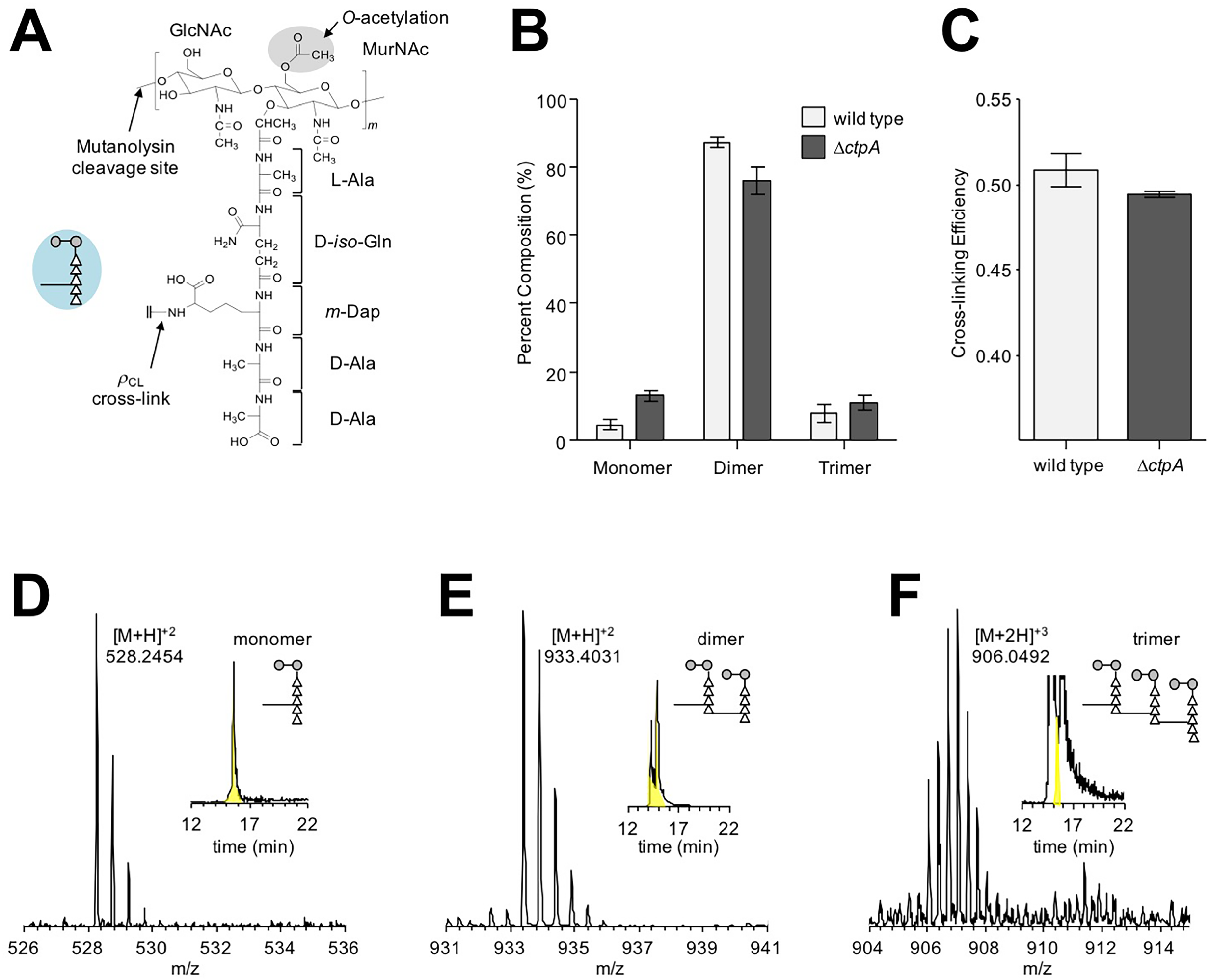
Changes in peptidoglycan composition of a Δ*ctpA* mutant. (A) Chemical structure of the peptidoglycan-repeat unit in *P. aeruginosa*. The disaccharide of the peptidoglycan-repeat unit is shown with two modifications: O-acetylation of MurNAc at C6 position and amidation of d-iso-Glu in the peptide stem to d-iso-Gln. The interpeptide bridge structure in *P. aeruginosa* is *m*-Dap. A schematic representation of a peptidoglycan-repeat unit is shown as a figure inset with the disaccharide as gray-filled circles, the amino acids in the pentapeptide stem as open triangles, and the interpeptide bridge structure as a line. The polymerized peptidoglycans are cross-liked by forming a peptide bond between the sidechain of *m*-Dap from one repeat unit to the carbonyl carbon of penultimate d-Ala of a peptidoglycan stem from a neighboring glycan chain. (B) The mutanolysin-digested muropeptide fragments were characterized by LC-MS. Forty-seven unique muropeptide ions from the wild type and thirty-three from the Δ*ctpA* mutant were identified from LC-MS and each muropeptide was quantified by integrating the extracted ion chromatogram (XIC) of the selected muropeptide ion. Peptidoglycan dimers are the most abundant muropeptides found in both wild type and Δ*ctpA* mutant with percent compositions of 87.30% ± 1.34% and 75.85% ± 3.90%, respectively. Muropeptide fragments larger than trimers were not observed. The *p*-values for the differences between wild type and Δ*ctpA* strains for monomers, dimers, and trimers based on the t-test are 0.0001, 0.0003, and 0.0216, respectively. (C) The calculated peptidoglycan cross-linking efficiency (*ρ_CL_*) for wild type and Δ*ctpA* mutant are 50.86% ± 0.98% and 49.46% ± 0.18%, respectively, with a *p*-value of 0.0038. All errors bars represent 95% confidence interval (n = 3). Representative mass spectra of monomer (D), dimer (E), and trimer (F) are shown with the XICs as insets. The corresponding chemical structures are in Supplementary Figure S1.

The cell walls of the Δ*ctpA* mutant had a reduced concentration of peptidoglycan dimers (75.85% ± 3.90%) compared to wild type (87.30% ± 1.34%), and a concomitant increase in monomers (Fig. 6B). This is consistent with decreased peptidoglycan cross-linking efficiency (*ρ_CL_*) in the Δ*ctpA* mutant. The calculated *ρ_CL_* for wild type and Δ*ctpA* mutant were 50.86% and 49.46%, respectively (Fig. 6C). The reduced cross-linking efficiency in the Δ*ctpA* mutant is small. However, the accompanying increase in trimers and monomers, indicates altered peptidoglycan biosynthesis. A small effect was expected because a *ctpA* mutant grows well, had few phenotypes in a Biolog Phenotype MicroArray (18), and is only slightly sensitive to low osmotic conditions (see below). Regardless, these data are consistent with the accumulation of peptidoglycan cross-link hydrolase activity in LbcA•CtpA-defective strains.

### LbcA enhances CtpA-dependent degradation of MepM *in vitro*

To test if MepM is a direct substrate of CtpA, we purified C-terminal hexahistidine tagged proteins. MepM was stable when incubated alone, but in the presence of CtpA the amount of MepM decreased (Fig. 7A). Furthermore, when MepM was incubated with both CtpA and LbcA it became undetectable, which is consistent with LbcA enhancing CtpA-dependent degradation (Fig. 7A). There was no MepM degradation when it was incubated with CtpA-S302A +/− LbcA. This supports the contentions that the S302A mutation destroys CtpA activity, that MepM degradation was due to CtpA and not a contaminating protease activity, and that LbcA cannot degrade MepM itself.

**FIG 7.**
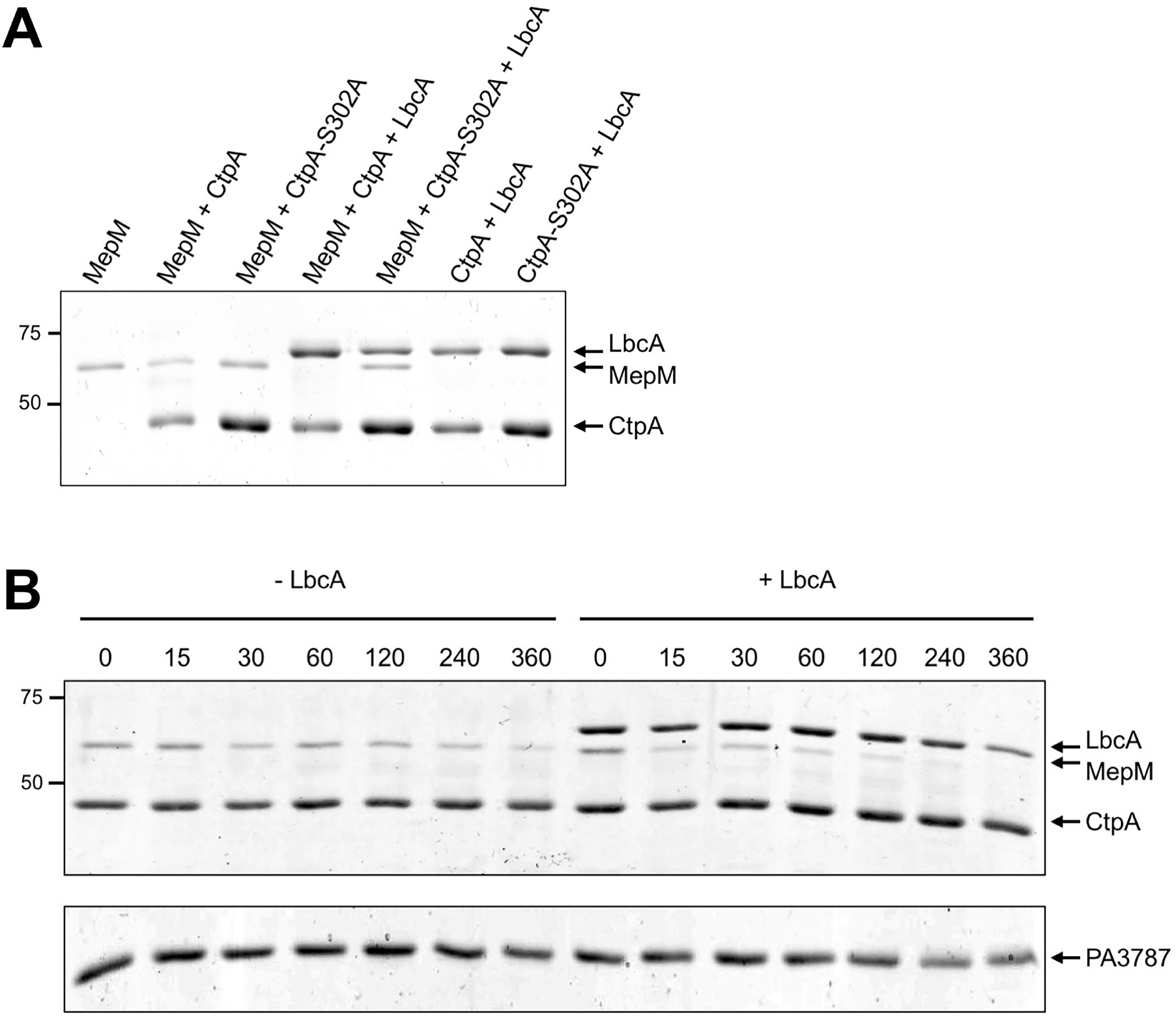
*In vitro* proteolysis of MepM. (A) CtpA degrades MepM and is enhanced by LbcA. 2 μM of the indicated C-terminal His_6_-tagged proteins were incubated for 3 h at 37°C. Samples were separated on a 12.5 % SDS PAGE gel, which was stained with coomassie brilliant blue. Approximate positions of molecular-mass-marker proteins (kDa) are indicated on the left-hand side. (B) Time course. Reactions contained 2 μM of CtpA-His_6_ and MepM-His_6_ either without (-LbcA) or with (+ LbcA) 2 μM of LbcA-His_6_. Reactions were terminated at the indicated time points by adding SDS-PAGE sample buffer and boiling. As a negative control, the same experiment was done using PA3787 in place of MepM. Samples were separated on 12.5 % SDS PAGE gels, which were stained with coomassie brilliant blue. Approximate positions of molecular-mass-marker proteins (kDa) are indicated on the left-hand side. PA3787 ran at the midway point between marker proteins of 37 and 25 kDa, which were above and below, respectively, the region of the gel shown in the figure.

We also monitored the CtpA-dependent degradation of MepM over time and found that it was faster in the presence of LbcA (Fig. 7B). Finally, we incubated CtpA with another LytM/M23 peptidase domain protein, PA3787. However, PA3787 was not degraded significantly, showing that CtpA has substrate specificity (Fig. 7B). These *in vitro* experiments support the characterization of MepM as a direct substrate of CtpA and suggest that LbcA catalyzes the proteolysis.

### The LbcA•CtpA complex degrades additional peptidoglycan cross-link hydrolases

MepM was the most abundant protein co-purified with LbcA•CtpA-S302A. Remarkably, the three proteins with the next highest PSMs are also known or predicted peptidoglycan hydrolases (PA1198, MltD and RlpA; Fig. 4B). PA1198 is homologous to *E. coli* NlpC/P60 peptidase family member MepS, a DD-endopeptidase that targets the same peptidoglycan cross-links as MepM (23). MltD and RlpA are lytic transglycosylases that attack the glycan chain (38–40).

To test if CtpA might cleave these putative peptidoglycan hydrolases, we constructed plasmids encoding arabinose inducible FLAG tagged versions. PA1198-FLAG was undetectable in the wild type, whereas it became abundant in a Δ*ctpA* mutant (Fig. 8A). We extended the PA1198 analysis to an Δ*lbcA* mutant, in which it also accumulated (Fig. 8A). Therefore, PA1198 might also be an LbcA•CtpA substrate. However, MltD-FLAG and RlpA-FLAG were detectible in wild type strains and did not accumulate in a Δ*ctpA* mutant (Fig. 8A). This suggests that MltD and RlpA are not CtpA substrates (see Discussion).

**FIG 8.**
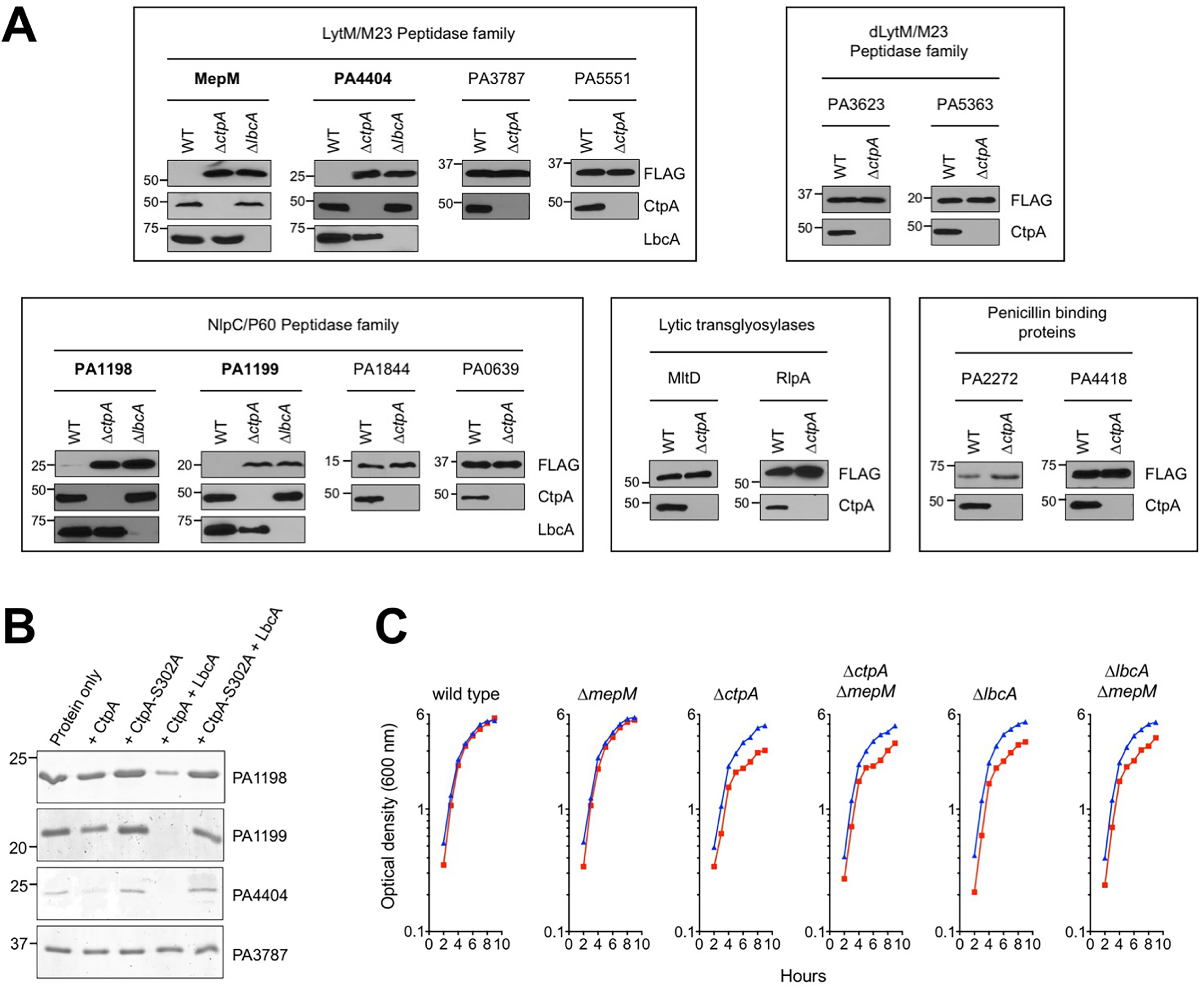
Evidence that the LbcA•CtpA complex has additional substrates. (A) *In vivo* analysis. Immunoblot analysis of equivalent amounts of whole cell lysates of wild type and Δ*ctpA* strains, and also of Δ*lbcA* strains for proteins that accumulated in the Δ*ctpA* mutant. All strains contained an arabinose-inducible expression plasmid encoding C-terminal FLAG-tagged versions of the indicated proteins. Approximate positions of molecular-mass-marker proteins (kDa) are indicated on the left-hand side. Test proteins were detected with anti-FLAG monoclonal antibodies, and CtpA and LbcA were detected with polyclonal antisera. Each box has analysis of proteins in related families, as indicated at the top. Only those proteins with names in bold face accumulated in the Δ*ctpA* mutant. (B) *In vitro* analysis. 2 μM of the indicated C-terminal His_6_-tagged proteins were incubated for 3 h at 37°C. Protein only indicates incubation of the test substrate alone. PA3787-His_6_ was used as a negative control. Samples were separated on a 15 % SDS PAGE gel, which was stained with coomassie brilliant blue. Approximate positions of molecular-mass-marker proteins (kDa) are indicated on the left-hand side. (C) Sensitivity of Δ*ctpA* and Δ*lbcA* mutants to low salt is not suppressed by Δ*mepM*. Strains were grown at 37°C in LB broth containing 1% (w/v) NaCl (blue) or no NaCl (red). Optical density was measured hourly.

We also used a biased approach by screening some putative *P. aeruginosa* peptidoglycan hydrolases. Again, this was done by constructing plasmids encoding arabinose inducible FLAG tagged versions. Candidates included three members of the LytM/M23 peptidase family that MepM is in (PA3787, PA4404 and PA5551), two members of the LytM/M23 family with predicted defective peptidase domains (PA3623/NlpD and PA5363), and three members of the NlpC/P60 family that PA1198 is in (PA0639, PA1199 and PA1844/Tse1). We also tested PA2272 (PBP3A) and PA4418 (PBP3), two homologs of *E. coli* penicillin binding protein 3 (PBP3/FtsI), which has been proposed to by proteolytically processed by its CTP, Prc/Tsp (8). Most of these candidates did not accumulate in a Δ*ctpA* mutant, suggesting that they are not substrates (Fig 8A). However, PA1199 and PA4404 were not detected in wild type but accumulated in the Δ*ctpA* mutant. We extended their analysis to an Δ*lbcA* mutant, in which they also accumulated (Fig. 8A). Therefore, PA1199 and PA4404 are also putative LbcA•CtpA proteolytic substrates.

To confirm PA1198, PA1199 and PA4404 as CtpA substrates, we purified hexahistidine tagged versions and tested them *in vitro*. All were degraded when incubated with LbcA and CtpA, but there was no degradation by CtpA-S302A (Fig. 8B). Therefore, CtpA degrades two peptidoglycan hydrolases in the LytM/M23 peptidase family (MepM and PA4404), and two in the NlpC/P60 family (PA1198 and PA1199). Their sequences predict all to be in the cell envelope, MepM as a peptidoglycan-associated protein and PA1198, PA1199 and PA4404 as outer membrane lipoproteins. Furthermore, all are predicted to cleave the peptide cross-links of peptidoglycan. Breaking these cross-links poses a danger to the bacterial cell and the LbcA•CtpA complex might be an important control mechanism used by *P. aeruginosa* to regulate this activity.

### LbcA•CtpA defective strains grow poorly in salt-free medium

*E. coli prc* mutants have altered cell morphology, and increased sensitivity to some drugs and low osmolarity (20). Biolog Phenotype Microarray analysis of a *P. aeruginosa* Δ*ctpA* mutant revealed no increased drug sensitivity (18). We have examined cells by light microscopy after growth at 37°C or 42°C, in media with or without salt, and in exponential and stationary phase, but Δ*ctpA* and Δ*lbcA* mutations did not noticeably affect cell shape or length (data not shown). This is not surprising because Prc and CtpA have different substrates (Prc processes FtsI, which might contribute to filamentation of Δ*prc* cells, whereas CtpA does not appear to process *P. aeruginosa* FtsI homologs, Fig. 8A). Also, *P. aeruginosa* has another CTP (PA3257/AlgO) that is a closer homolog of *E. coli* Prc than is CtpA. Even so, if multiple peptidoglycan hydrolases accumulate in Δ*ctpA*/Δ*lbcA* mutants (Fig. 8A), and peptidoglycan cross-linking is compromised (Fig. 6), then sensitivity to low osmolarity is expected. Therefore, we also investigated this possibility.

Δ*ctpA* and Δ*lbcA* mutants grew more slowly than wild type in LB broth without NaCl (Fig. 8C). This phenotype was not exacerbated at 42°C and/or on agar (data not shown). Regardless, the occurrence and subtle nature of this phenotype is consistent with the slightly reduced peptidoglycan cross-linking efficiency (Fig. 6). Importantly, this phenotype was not fully suppressed by a Δ*mepM* mutation, which further supports LbcA•CtpA degrading multiple peptidoglycan hydrolases.

## DISCUSSION

We have discovered that *P. aeruginosa* CtpA forms a complex with LbcA, and both proteins are required for CtpA-mediated proteolysis *in vivo* and *in vitro*. We identified four proteolytic substrates, all of which are predicted to cleave the cross-links between peptidoglycan chains. Therefore, the LbcA•CtpA complex might be an important control mechanism used by *P. aeruginosa* to regulate peptidoglycan cross-link hydrolysis. It has been difficult to investigate if substrate degradation is regulated, because they are difficult to detect in protease-competent strains (e.g. Figs. 4 and 8). However, even though CtpA and LbcA protein levels appear constant regardless of growth stage, our preliminary evidence suggests that at least one substrate might be degraded at different rates (D. Srivastava and A. J. Darwin, unpublished data). Investigating how this occurs, and the signal(s) involved, are intriguing future questions.

An exciting parallel exists between the complex we have discovered, and the NlpI•Prc complex in *E. coli*. Prc is a CTP, and NlpI is an outer membrane lipoprotein with TPR motifs, and together they degrade one peptidoglycan hydrolase, MepS (22). The similarities to LbcA•CtpA are obvious. However, Prc and CtpA are evolutionarily divergent, differing in size by approximately 30 kDa and belonging to different CTP subfamilies. Although NlpI and LbcA are both outer membrane lipoproteins containing short TPR motifs, their primary sequences are not similar and LbcA is much larger than NlpI. In the *E. coli* NlpI•Prc complex, an NlpI homodimer binds two Prc monomers to create docking sites for the MepS substrate (41). The LbcA•CtpA complex in *P. aeruginosa* could have similarities to this, but we suspect that its stoichiometry and functioning will have differences, due to the divergence of the proteins involved. Regardless, it is important that two very different CTPs, and two very different lipoproteins, form complexes in two divergent bacterial species to degrade peptidoglycan hydrolases. This suggests that this could be a widespread phenomenon and a new paradigm might emerge in which many bacterial CTPs work with partner proteins to target cell wall hydrolases.

LbcA•CtpA degrades four predicted peptidoglycan cross-link hydrolases (Figs. 4, 7-9). Consistent with this, a Δ*ctpA* mutant has a cell envelope structural defect when viewed by electron microscopy (18), peptidoglycan cross-linking is reduced (Fig. 6), and it grows poorly in LB broth without NaCl (Fig. 8C). All of this suggests a weakened cell wall caused by increased peptidoglycan cross-link hydrolase activity. Removing the MepM substrate did not suppress the slow growth of Δ*ctpA* and Δ*lbcA* mutants in salt free medium (Fig. 8C). This indicates that at least one other CtpA substrate must contribute to this phenotype. In fact, although it is not obvious from the growth curves, the Δ*mepM* mutation did improve growth very slightly. Therefore, we hypothesize that the poor growth in LB broth without NaCl results from the combined effect on the cell wall of two or more accumulating CtpA substrates. We have not yet been able to make null mutants of the other CtpA substrates, but a future goal is to investigate their roles.

The defective T3SS and accelerated surface attachment phenotypes of a Δ*ctpA* mutant are most likely to be secondary consequences of a compromised cell wall. However, these phenotypes were efficiently suppressed by a Δ*mepM* mutation. MepM was the most abundant protein trapped by the LbcA•CtpA-S302A complex (Fig. 4B) and so it might be having the largest influence on the cell wall. If so, even though removing MepM might not fix all the cell wall problems, it might improve things sufficiently to prevent effects on the T3SS and surface attachment. However, another possibility is that MepM plays a more specialized role. In this regard it is intriguing to note that the assembly of transenvelope systems, including a T3SS, requires rearrangement of the peptidoglycan (42–44). Some T3SS-encoding loci encode a peptidoglycan hydrolase that is needed for T3SS function (45, 46). Unlike MepM, these are lytic transglycosylases, but it is still possible that a specific role of MepM is important for T3SS assembly or function in *P. aeruginosa*. Indeed, many T3SS loci do not encode a dedicated peptidoglycan hydrolase, raising the possibility that they rely on endogenous cellular hydrolase(s). It is also worth noting the possibility that some peptidoglycan hydrolases function in a complex and removing one of them destroys the function of the entire complex. The lytic transglycosylases MltD and RlpA were pulled down by the LbcA•CtpA-S302A trap, but they do not appear to be CtpA substrates (Figs 4B and 8A). This could be explained by an interaction with a CtpA substrate such as MepM.

The MepM and PA4404 substrates are in the LytM/M23 family of peptidases, and both have homology to *E. coli* MepM. However, indirect genetic evidence suggests that Prc, the only CTP in *E. coli*, might not degrade MepM in that species (47). *E. coli* Prc does degrade the NlpC/P60 peptidase family member MepS (22). This is notable, because the CtpA substrates PA1198 and PA1199 are both homologous to *E. coli* MepS. Therefore, despite being from divergent CTP subfamilies, *P. aeruginosa* CtpA and *E. coli* Prc both degrade MepS-like hydrolases. This is surprising because *P. aeruginosa* has a second CTP, and the other one is a homolog of *E. coli* Prc (PA3257, also known as AlgO). The abundance of PA1198 and PA1199 in a Δ*ctpA* strain (Fig. 8A) suggests that Prc might not degrade MepS hydrolases in *P. aeruginosa*.

*P. aeruginosa* Prc was proposed to cleave the antisigma factor MucA, inducing the extracytoplasmic function (ECF) sigma factor AlgT/U and alginate production (26–28). However, the AlgT/U regulon can also be induced by cell wall-inhibitory antibiotics (28). Therefore, it is possible that *P. aeruginosa* Prc cleaves one or more peptidoglycan modifying enzymes, and its impact on the AlgT/U regulon is a consequence of altered peptidoglycan rather than Prc cleaving MucA directly. In fact, CtpA overproduction induces an ECF sigma factor that can be activated by the cell wall-inhibitory antibiotic D-cycloserine, but CtpA is unlikely to cleave its antisigma factor (18). Prc has also been implicated in activating several other ECF sigma factors in *Pseudomonas* species (48). However, induction of ECF sigma factors by cell wall stress is common, and an effect of Prc on peptidoglycan could perhaps be the explanation. Importantly, Prc has not been shown to cleave any antisigma factor directly.

In summary, a binary protein complex degrades at least four proteins predicted to cleave peptidoglycan cross-links in *P. aeruginosa* (Fig. 9). Therefore, LbcA•CtpA might be an important mechanism used to control peptidoglycan hydrolysis, a reaction with the potential for catastrophic consequences if it is not carefully constrained. Importantly, similarities to the distantly related NlpI•Prc system of *E. coli* suggest that this might be a widespread phenomenon in diverse bacteria.

**FIG 9.**
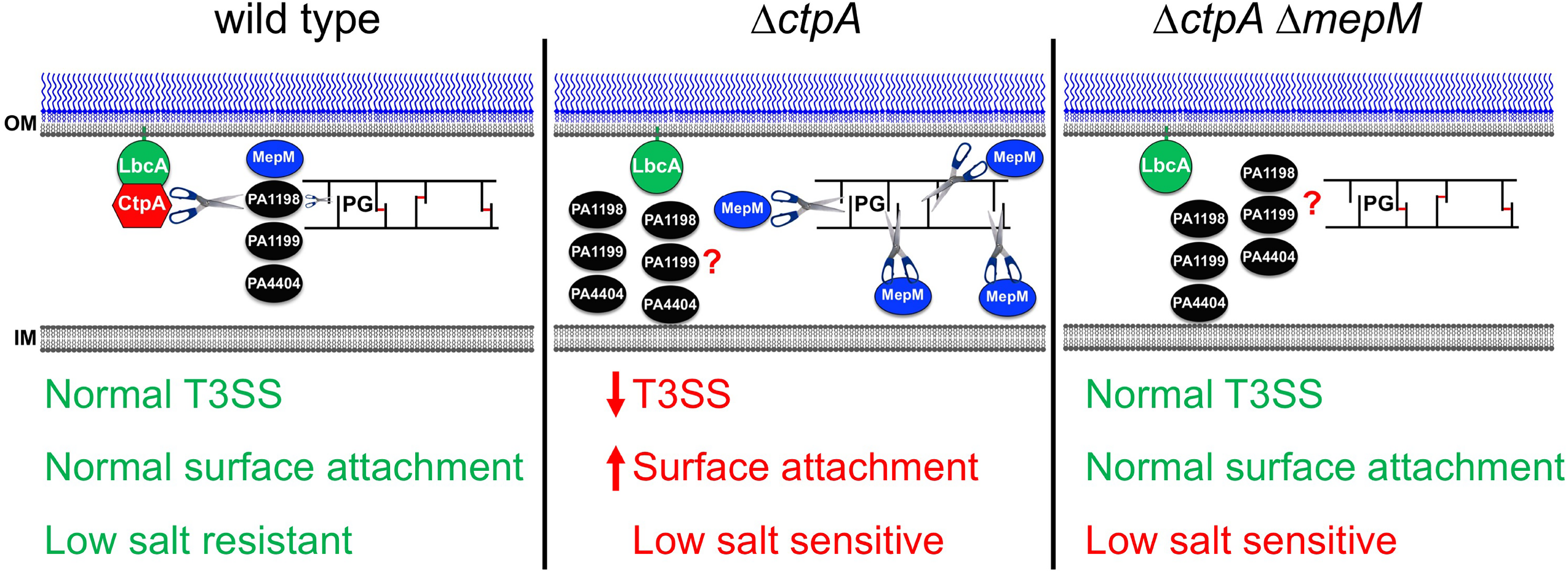
Summary. In a wild type strain, CtpA forms a complex with the outer membrane lipoprotein LbcA, which is required for CtpA to cleave its targets. The LbcA•CtpA complex can degrade at least four enzymes predicted to cleave peptidoglycan cross-links (MepM, PA1198, PA1199 and PA4404). This might provide a mechanism to carefully constrain crosslink hydrolysis, limiting it to the amount needed for roles such as cell elongation and/or division. In a Δ*ctpA* mutant, the peptidoglycan hydrolases can accumulate, which causes excessive crosslink cleavage and subsequent phenotypes, including a defective T3SS, enhanced surface attachment and low salt sensitivity. When *mepM* is deleted from the Δ*ctpA* mutant, the T3SS and surface attachment phenotypes revert back to wild type, suggesting that elevated MepM activity was their primary cause. However, low salt sensitivity of a Δ*ctpA* mutant is not suppressed by Δ*mepM*, suggesting that increased activity of one or more of the other substrates is also compromising cell wall integrity. OM = outer membrane, IM = inner membrane.

## MATERIALS AND METHODS

### Bacterial strains and standard growth conditions

Strains and plasmids are listed in Table 1. Bacteria were grown routinely in Luria-Bertani (LB) broth or on LB agar at 37°C. During conjugation procedures, *P aeruginosa* was occasionally selected on Vogel-Bonner minimal (VBM) agar. *E. coli* K-12 strain SM10 was used to conjugate plasmids into *P. aeruginosa* (49).

**TABLE 1.**
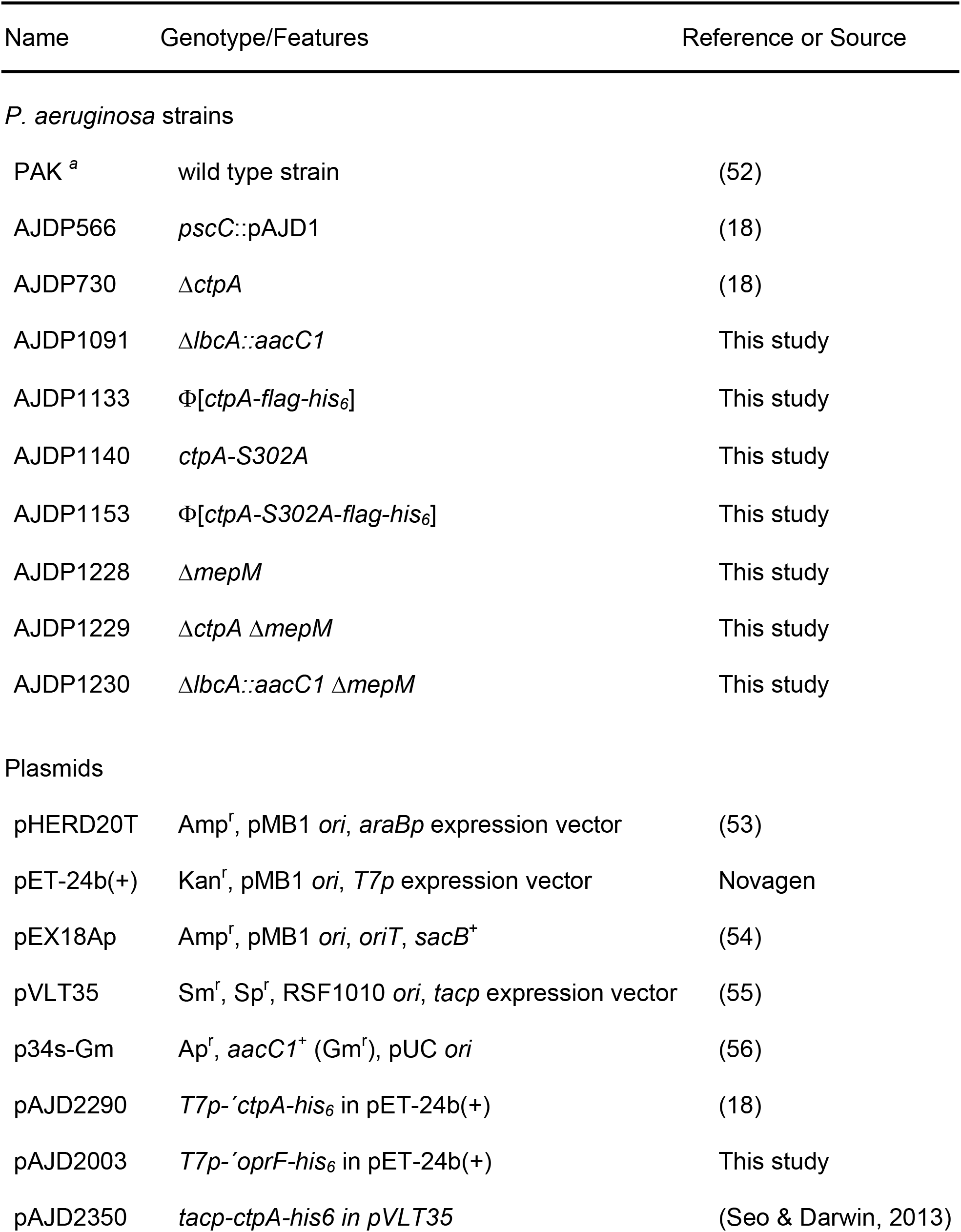
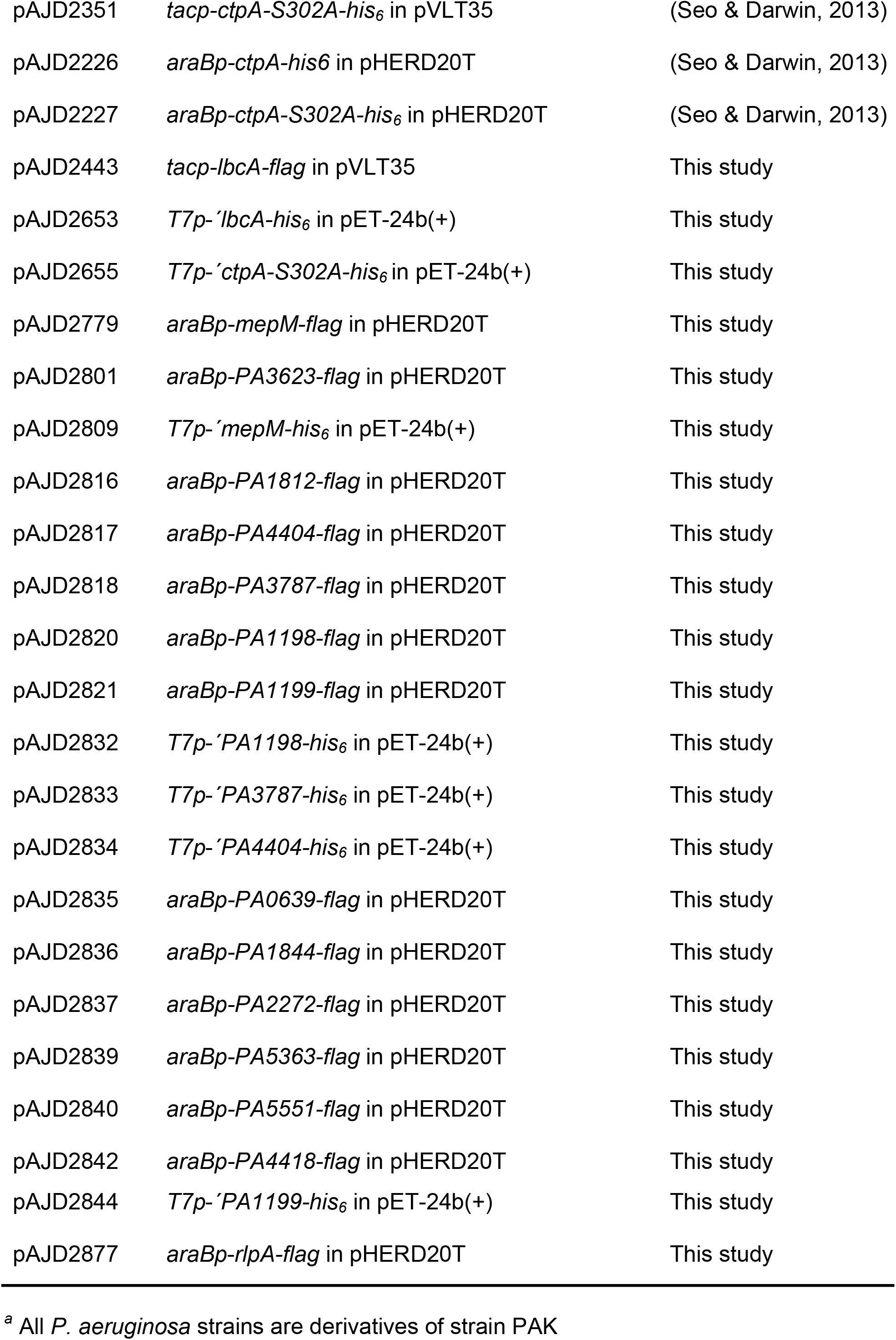
Strains and plasmids

| Name | Genotype/Features | Reference or Source |
| --- | --- | --- |
| *P. aeruginosa* strains |  |  |
| PAK <sup>a</sup> | wild type strain | (52) |
| AJDP566 | *pscC::pAJD1* | (18) |
| AJDP730 | Δ*ctpA* | (18) |
| AJDP1091 | Δ*lbcA::aacC1* | This study |
| AJDP1133 | $\Phi$[*ctpA-flag-his<sub>6</sub>*] | This study |
| AJDP1140 | *ctpA-S302A* | This study |
| AJDP1153 | $\Phi$[*ctpA-S302A-flag-his<sub>6</sub>*] | This study |
| AJDP1228 | Δ*mepM* | This study |
| AJDP1229 | Δ*ctpA* Δ*mepM* | This study |
| AJDP1230 | Δ*lbcA::aacC1* Δ*mepM* | This study |
| Plasmids |  |  |
| pHERD20T | Amp<sup>r</sup>, pMB1 *ori*, *araBp* expression vector | (53) |
| pET-24b(+) | Kan<sup>r</sup>, pMB1 *ori*, *T7p* expression vector | Novagen |
| pEX18Ap | Amp<sup>r</sup>, pMB1 *ori*, *oriT*, *sacB*<sup>+</sup> | (54) |
| pVLT35 | Sm<sup>r</sup>, Sp<sup>r</sup>, RSF1010 *ori*, *tacp* expression vector | (55) |
| p34s-Gm | Ap<sup>r</sup>, *aacC1*<sup>+</sup> (Gm<sup>r</sup>), pUC *ori* | (56) |
| pAJD2290 | *T7p-ctpA-his<sub>6</sub>* in pET-24b(+) | (18) |
| pAJD2003 | *T7p-oprF-his<sub>6</sub>* in pET-24b(+) | This study |
| pAJD2350 | *tacp-ctpA-his<sub>6</sub>* in pVLT35 | (Seo & Darwin, 2013) |

|  |  |  |  |
| --- | --- | --- | --- |
| 780 | pAJD2351 | *tacp-ctpA-S302A-his<sub>6</sub>* in pVLT35 | (Seo & Darwin, 2013) |
| 781 | pAJD2226 | *araBp-ctpA-his<sub>6</sub>* in pHERD20T | (Seo & Darwin, 2013) |
| 782 | pAJD2227 | *araBp-ctpA-S302A-his<sub>6</sub>* in pHERD20T | (Seo & Darwin, 2013) |
| 783 | pAJD2443 | *tacp-lbcA-flag* in pVLT35 | This study |
| 784 | pAJD2653 | *T7p-<sup>-</sup>lbcA-his<sub>6</sub>* in pET-24b(+) | This study |
| 785 | pAJD2655 | *T7p-<sup>-</sup>ctpA-S302A-his<sub>6</sub>* in pET-24b(+) | This study |
| 786 | pAJD2779 | *araBp-mepM-flag* in pHERD20T | This study |
| 787 | pAJD2801 | *araBp-PA3623-flag* in pHERD20T | This study |
| 788 | pAJD2809 | *T7p-<sup>-</sup>mepM-his<sub>6</sub>* in pET-24b(+) | This study |
| 789 | pAJD2816 | *araBp-PA1812-flag* in pHERD20T | This study |
| 790 | pAJD2817 | *araBp-PA4404-flag* in pHERD20T | This study |
| 791 | pAJD2818 | *araBp-PA3787-flag* in pHERD20T | This study |
| 792 | pAJD2820 | *araBp-PA1198-flag* in pHERD20T | This study |
| 793 | pAJD2821 | *araBp-PA1199-flag* in pHERD20T | This study |
| 794 | pAJD2832 | *T7p-<sup>-</sup>PA1198-his<sub>6</sub>* in pET-24b(+) | This study |
| 795 | pAJD2833 | *T7p-<sup>-</sup>PA3787-his<sub>6</sub>* in pET-24b(+) | This study |
| 796 | pAJD2834 | *T7p-<sup>-</sup>PA4404-his<sub>6</sub>* in pET-24b(+) | This study |
| 797 | pAJD2835 | *araBp-PA0639-flag* in pHERD20T | This study |
| 798 | pAJD2836 | *araBp-PA1844-flag* in pHERD20T | This study |
| 799 | pAJD2837 | *araBp-PA2272-flag* in pHERD20T | This study |
| 800 | pAJD2839 | *araBp-PA5363-flag* in pHERD20T | This study |
| 801 | pAJD2840 | *araBp-PA5551-flag* in pHERD20T | This study |
| 802 | pAJD2842 | *araBp-PA4418-flag* in pHERD20T | This study |
| 803 | pAJD2844 | *T7p-PA1199-his<sub>6</sub>* in pET-24b(+) | This study |
| 804 | pAJD2877 | *araBp-rlpA-flag* in pHERD20T | This study |

<sup>a</sup> All *P. aeruginosa* strains are derivatives of strain PAK

### Plasmid and strain constructions

To construct in frame deletion mutants, two fragments of ~ 0.5 kb each corresponding to regions upstream and downstream of the deletion site were amplified by PCR and cloned into pEX18Ap. The plasmids were integrated into the *P. aeruginosa* chromosome after conjugation from *E. coli* and sucrose resistant, carbenicillin sensitive segregants were isolated on LB agar containing 10% sucrose. Deletions were verified by colony PCR. Δ*lbcA::aacC1* mutants were constructed similarly, except the *aacC1* cassette from plasmid p34s-Gm was cloned between the upstream and downstream fragments in pEX18Ap.

To make strain AJDP1140 encoding CtpA-S302A, two ~ 0.5 kb fragments flanking codon 302 of *ctpA* were amplified by PCR chromosome. For each fragment one of the primers incorporated a mismatch at codon 302 to convert it to encode alanine. The fragments were joined in a PCR SOEing reaction (50) via their overlapping regions around codon 302. The product was cloned into pEX18Ap and exchanged for the corresponding region of the *ctpA* gene by integration, selection for sucrose-resistant segregants and confirmation by colony PCR and DNA-sequencing.

To make strains encoding CtpA-FLAG-His_6_ and CtpA-S302A-FLAG-His_6_, two ~ 500 bp fragments surrounding the stop codon of *ctpA* were amplified by PCR. For each fragment one primer incorporated a region encoding FLAG-His_6_. The fragments were joined in a PCR SOEing reaction via their overlapping FLAG-His_6_ sequences. The products were cloned into plasmid pEX18Ap and used to fuse the FLAG-His_6_ encoding region to the *ctpA* gene by integration, selection for sucrose-resistant segregants and confirmation by colony PCR and DNA-sequencing.

Plasmids encoding C-terminal FLAG-tagged proteins were constructed by amplifying genes using a downstream primer that incorporated a region encoding the FLAG tag, and then cloning into pHERD20T or pVLT35. Plasmids for protein overproduction and purification were constructed by PCR amplification of genes, without their predicted N-terminal signal sequences and stop codons, and cloning into pET-24b(+) to encode C-terminal His_6_-tagged versions.

### Polyconal antiserum production and immunoblotting

*E. coli* strain ER2566 (NEB) containing a pET-24b(+) derivative encoding MepM-His_6_, LbcA-His_6_ or OprF-His_6_ was grown in LB broth to mid-log phase at 37°C with aeration. Protein production was induced with 1mM IPTG for 4 hrs at 37°C for LbcA-His_6_ and OprF-His_6_, or overnight at 16°C for MepM-His_6_. Proteins were purified under denaturing conditions by nickel-nitrilotriacetic acid (NTA) affinity chromatography as described by the manufacturer (Qiagen). Polyclonal rabbit antisera were raised by Covance Research Products Inc.

Semi dry electroblotting was used to transfer SDS-PAGE separated samples to nitrocellulose. Chemiluminescent detection followed incubation with polyclonal antiserum or monoclonal antibody, then goat anti-rabbit IgG (Sigma) or goat anti-mouse IgG (Sigma) horseradish peroxidase conjugates used at the manufacturers recommended dilution. Primary antisera or antibodies were diluted 5,000-fold for anti-THE-His (GenScript) and anti-FLAG M2 (Sigma), 7,500-fold for anti-MepM, 8,000-fold for anti-LbcA, 10,000-fold for anti-PA0943, anti-PA4068 and anti-CtpA (18, 51) and 20,000 fold for anti-OprF.

### CtpA-FLAG-His_6_ tandem purification

Strains were grown in 100 ml LB broth at 37°C with aeration to OD 600 nm ~ 1.2 - 1.4 and cells were collected by centrifugation. For formaldehyde cross-linking, the pellet was washed with cold 10 mM potassium phosphate buffer pH 8.0 and resuspended to an OD 600 nm of 5 in the same buffer. 1% formaldehyde was added followed by incubation at room temperature for 30 min. 0.3 M Tris-HCl pH 7.5 was added to quench and the cells were collected by centrifugation. Pellets were resuspended in 3 ml lysis buffer (50 mM Tris-HCl pH 7.5, 150 mM NaCl, 5 mM Imidazole), and Roche complete protease inhibitors (2x concentration), 1 μg/ml DNaseI, 1 μg/ml RNase, and 1 mg/ml lysozyme were added. Cells were disrupted by sonication, then 1% *n*-dodecyl β-D-maltoside (DDM) was added followed by incubation with rotation for 30 min at 4°C. Insoluble material was removed by centrifugation at 13,000 × *g* for 30 min at 4°C. 200 μl of nickel-NTA agarose in lysis buffer was added to the supernatant, followed by incubation with rotation for 1.5 h at 4°C. The resin was collected in a drip column and washed with 4 ml lysis buffer, then 4 ml lysis buffer containing 20 mM imidazole. Proteins were eluted in 1 ml lysis buffer containing 250 mM imidazole, mixed with 30 μl anti-FLAG M2 agarose resin (Sigma) in TBS (10 mM Tris-HCl pH 7.5, 150 mM NaCl), and incubated with rotation for 2 h at 4°C. A 1 ml spin column (Pierce 69725) was used to wash the resin three times with 800 μl TBS. Proteins were eluted by adding 100 μl of 2 μg/ml FLAG peptide (F3290 Sigma) in TBS and incubating with rotation at 4°C for 30 min. Proteins were identified by LC-mass spectrometry (NYU School of Medicine Proteomics Laboratory).

### CtpA-His_6_, LbcA-FLAG tandem purification

Δ*ctpA* strains containing pVLT35 derivatives encoding CtpA-His_6_ or CtpA-S302A-His_6_, and pHERD20T encoding LbcA-FLAG, were inoculated to an OD 600 nm of 0.05 in 250 ml LB broth containing 5 mM EGTA, 100 μM IPTG and 0.02% (w/v) arabinose. Cultures were incubated at 37°C with 225 rpm rotation until the OD 600 nm was ~ 1.5. Procedures described in the preceding section were used for protein purification and identification.

### Analysis of T3SS substrates in cell free supernatants

Strains were inoculated at OD 600 nm of 0.04 in test tubes containing 5 ml Tryptic Soy Broth (TSB) containing 5 mM EGTA, and grown for 6 h at 37°C on a roller. Cells from the equivalent of 1 ml of culture at OD 600 nm of 1.5 were removed by centrifugation at 3,300 × *g* for 15 min at 4°C. The supernatant was filtered with a 0.22 μm filter, then 1/10^th^ volume trichloroacetic acid was added, followed by incubation at 4°C overnight. Proteins were collected by centrifugation at 4,500 × *g* for 40 min at 4°C, washed twice with 1 ml ice cold acetone, dried at room temperature for 30-40 min, and resuspended in SDS-PAGE sample buffer. Proteins were separated by SDS PAGE and stained with silver.

### Surface attachment

Saturated cultures in LB broth were diluted to an OD 600 nm of 0.1 in 13 × 100 mm borosilicate glass tubes containing 0.5 ml of M63 salts supplemented with 1 mM MgSO_4_, 0.5% (w/v) casamino acids and 0.2% (w/v) glucose. The tubes were incubated at 30°C without agitation for 8 h. Culture medium was removed and the tubes were washed twice with 2 ml of water. 500 μl of 0.1% (w/v) crystal violet was added followed by incubation at room temperature for 10 min, two washes with 10 ml water, and then the tubes were inverted for 10 min at room temperature and photographed. The crystal violet was dissolved in 1 ml ethanol and absorbance at 595 nm was measured. A tube that contained growth medium alone was used to generate the blank.

### Subcellular fractionation

Osmotic shock was done as previously (18), except that cultures were grown to OD 600 nm of ~ 1. To generate soluble and insoluble fractions, 100 ml cultures in LB broth were grown at 37°C with shaking at 225 rpm to an OD 600 nm of ~ 1. Cells were collected by centrifugation, washed with 10 ml of 20 mM Tris-HCl pH 7.5, 20 mM EDTA (TE), resuspended in 10 ml cold TE containing Roche complete protease inhibitors, and lysed by two passages through a French pressure cell (1,100 psi). Intact cells were removed by centrifugation at 8,000 × *g* for 10 min at 4°C. The supernatant was centrifuged at 100,000 × *g* for 1 h at 4°C. The supernatant was the soluble fraction. The pellet (membrane fraction) was washed twice with 20 mM Tris-HCl pH 7.5, and resuspended in 20 mM Tris-HCl pH 7.5 containing Roche complete protease inhibitors.

### CHO-K1 cell cytotoxicity

Chinese hamster ovary cells were grown in Roswell Park Memorial Institute (RPMI) 1640 medium with 10% fetal bovine serum (FBS) and 2 mM glutamine, at 37°C in a 5% CO_2_ atmosphere. 2×10^5^ cells/well were seeded in a 24-well plate and incubated at 37°C for 16 h. Attached cells were washed with phosphate buffered saline (PBS) and covered with RPMI 1640 containing 1% FBS and 2 mM glutamine. *P. aeruginosa* was grown in TSB at 37°C until the OD 600 nm was 0.8, then added to the CHO-K1 cells at a multiplicity of infection of ~ 10. The plate was centrifuged at 250 × *g* for 5 min and incubated for 4 h at 37°C. Supernatants were collected after centrifugation at 250 × *g* for 5 min and lactate dehydrogenase (LDH) concentration was measured with the LDH Cytotox assay (Promega).

### Cell wall isolation and digestion

Bacteria were grown to saturation in 500 ml LB broth at 37 °C with 150 rpm shaking. Cells were harvested by centrifuging at 5,200 × *g* for 20 min, resuspended in PBS, and sterilized by boiling for 60 min. Samples were bead beaten (Bead Mill 24; Fisher Scientific) with 0.5 mm diameter glass beads for ten 1 min cycles with 1 min rests. Beads were removed using a Steriflip 20 μm nylon vacuum filter (EMD Millipore). Crude cell wall pellets were resuspended in 2 ml PBS, to which 8 ml of 2% SDS (w/v) was added. They were boiled for 60 min, washed with deionized water, and resuspended in 2 ml of 50 mM Tris pH 8.0. 200 μg DNase was added and incubated at 37°C for 24 h and 150 rpm, followed by 200 μg pronase E and another 24 h incubation. Cell walls were washed once and resuspended in 1 ml of 50 mM Tris pH 8.0, then digested by adding 0.5 KU of mutanolysin (Sigma-Aldrich) and incubating at room temperature for 24 h. Another 0.5 KU of mutanolysin was added followed by another 24 h incubation. Digested cell walls were frozen, lyophilized, and dissolved in 1 ml of 0.375 M sodium borate buffer (pH 9.0) using HPLC-grade water. Muropeptides were reduced by adding 10 mg of sodium borohydride at room temperature for 30 min. Reduction was quenched by adding 125 μl phosphoric acid. Samples were frozen at - 80°C, lyophilized, resuspended in 1 ml of 1% trifluroacetic acid, centrifuge filtered, and cleaned up for LC-MS using 100 μl Pierce C18 tips.

### Peptidoglycan analysis by liquid chromatography-mass spectrometry

Mutanolysin-digested muropeptide fragments were separated using a NanoACQUITY Ultra Performance Liquid Chromatography System (Waters). A reverse phase BEH C18 column (length 100 mm, diameter 75 μm) of 1.7 μm bead with pore size of 130 Å was used. Separation was carried out by injecting 2 μl of mutanolysin-digested PG from a 5 μl sample loop to the column under isocratic condition of 98% mobile phase A (0.1% formic acid in HPLC water) and 2% mobile phase B (90% acetonitrile & 10% of HPLC water added with 0.1% formic acid) for 5 min, then a linear gradient to 50% buffer B was applied for 30 min. The column was regenerated under isocratic condition with 85% buffer B for 5 min, a linear gradient to 98% buffer A for 1 min, then isocratic at 98% buffer A for 23 min. The flow rate was kept constant (0.6 μl/min) throughout the analysis. Fibrinopeptide B (Glu-Fib) was used as an internal standard. Data were analyzed using MassLynx (Waters) and MATLAB (MathWorks).

### *In vitro* proteolysis

*E. coli* ER2566 (NEB) containing each pET-24b(+) derivative were grown in 1L LB broth at 37°C with aeration until the OD 600 nm was 0.6-0.8. Protein production was induced by adding 1 mM IPTG and incubating for 16 h at 16°C with aeration. Proteins were purified under native conditions by NTA agarose affinity chromatography as recommended by the manufacturer (Qiagen). Following elution with imidazole, proteins were exchanged into 50 mM NaH_2_PO_4_, 300 mM NaCl pH 8.0, using Amicon Ultra-4 centrifuge filter devices (10 kDa cutoff), then supplemented with 10% glycerol and stored at −20°C. However, CtpA-His_6_ and CtpA-S302A-His_6_ only were eluted sequentially with 50 mM NaH_2_PO_4_, 300 mM NaCl, pH 8.0, containing 50 mM, 100 mM, 150 mM, 200 mM or 250 mM immidazole. Buffer exchange and freezing was not possible due to precipitation. Therefore, CtpA-His_6_ and CtpA-S302A-His_6_ were purified, stored at 4°C, and used within 48 h. All experiments were done at least three times with representative experiments shown in the figures.

### Growth curves

Saturated cultures were diluted into 5 ml of LB broth, containing either 1% (w/v) NaCl or no NaCl, in 18 mm diameter test tubes so that the initial OD 600 nm was approximately 0.1. The cultures were grown on a roller drum at 37°C for 9 h and 0.1 ml samples were removed at hourly intervals for OD 600 nm measurement.

## ACKNOWLEDGEMENTS

We thank Sindhoora Singh for constructing plasmid pAJD2003. Research was supported by the National Institute of Allergy and Infectious Diseases (NIAID) of the National Institutes of Health, under Award Numbers R21AI117131 and R01AI136901. The content is solely the responsibility of the authors and does not necessarily represent the official views of the National Institutes of Health. Mass spectrometric protein identification done by NYU School of Medicine’s proteomics laboratory was partly supported by NYU School of Medicine. S.J.K. and B.R. conducted the peptidoglycan analysis and analyzed the data, were supported by the National Institutes of Health Grant GM116130, and acknowledge the Baylor University Mass Spectrometry Centre (BU-MSC) for support.

---

**Supplemental Table 1**. Proteins that co-purified in each of two LbcA•CtpA-S302A pulldowns, but were absent in each of two LbcA•CtpA pulldowns

**Supplemental Figure S1**. Chemical structure of peptidoglycan monomer, dimer, and trimer corresponding to the mass spectra shown in Figure 6

